# Whole-genome sequencing of a mid-20^th^-century femur from central Israel in an open missing-person case

**DOI:** 10.64898/2026.04.24.720291

**Authors:** Ethel Vol, Shamam Waldman, Ariel Lomes, Esther S. Brielle, Noam Appel, Boris Dolin, Sharon Asif, Yossi Nagar, Eyal Marco, Nathaniel Bergman, Oded Khaner, Dvir Raviv, Jacob Oliel, Rafael Y. Lewis, Shai Carmi

## Abstract

Genome-wide technologies can generate investigative leads in cold cases by determining the genetic ancestry of the forensic sample. Increasingly, DNA extraction and whole-genome sequencing or genotyping are being used to analyze early or middle-20^th^ century skeletal remains. Here, we present the first case, to our knowledge, of whole-genome sequencing of a middle-20^th^-century bone sample from the Middle East. A femur discovered in a cave in Central Israel was proposed to belong to a person of Ashkenazi Jewish ancestry who was missing since 1948. Following DNA extraction and single-stranded library preparation, whole-genome sequencing generated nearly 500 million reads. However, only 0.5% of the reads mapped to the human genome, providing depth of coverage of 0.07x. After quality control and male sex inference, ancestry assignment was performed using principal components and ADMIXTURE analyses. The results suggested that the genome definitively belonged to a person of Arab ancestry, refuting the hypothesis of an Ashkenazi Jewish origin.

## Introduction

Next generation sequencing and genotyping technologies have revolutionized forensic investigations, permitting the simultaneous and high-resolution determination of uniparental haplogroups, genetic ancestry, and genetic relatedness^1,2^. Recent applications of genome-wide methods in forensic genetics have focused on samples from the past few decades, often with the goal of pursuing a forensic genetic genealogy search for distant living relatives^3,4^. Experimental and computational innovations from ancient DNA academic research can expedite the generation and analysis of genome-wide data for even older forensic samples, potentially leading to high-resolution ancestry assignment and detection of relatives^5–7^.

Accordingly, several recent studies have utilized genome-wide technologies, including whole-genome sequencing and targeted genotyping, to study forensic samples from the early/mid-20^th^ century. These studies included a 1916 Irish rebel^8^, soldiers killed in the Korea War and World War II^9^, a family member of the last Russian Tzar^10^, World War II victims in Greece^11^, Spanish Civil War victims^12^, an Idaho homicide victim^13^, an Australian drowning victim^14^, and a Belgian mining disaster victim^15^, among other cases^16^. However, no study to our knowledge has yet attempted whole-genome sequencing of early/mid-20^th^ century forensic samples from the Middle East, even though many human remains from that period are still unidentified. Genome-wide data could thus provide crucial forensic leads.

### Case description

A search for the remains of four Israeli persons, missing since 1948, was conducted a few years ago in the Shephelah region in central Israel. The search identified the skeletal remains of a single human individual in a cave near the place where these persons were last seen.

The remains were characterized using multiple approaches. The skeleton was partially articulated, suggesting a primary burial that was disturbed. The skeleton was put on top of the cave’s sand deposits, and its upper surface showed some signs of exposure to fire. The skull and pelvis manifested male morphology, as the glabella and the eyebrows, as well as the mastoid process and supra-mastoid crest, were markedly developed, and the pelvic subpubic angle was relatively narrow^20^. The age at death was estimated as 25-30 years, based on a combination of skeletal markers such as suture closure in the long bones, the vertebrae, and the maxillary hard palate^21–23^; tooth attrition rate^24^; fusion of the sagittal suture^25^; and chronological changes in the pelvic symphysis and the sternal end of ribs^26,27^. A femur and a rib from the skeleton were radiocarbon dated at the Dangoor Research Accelerator Mass Spectrometer laboratory of the Weizmann Institute. The results suggested a date of death in the early-middle 20^th^ century (in addition to a few earlier possible periods).

DNA was extracted from the remains and an STR profile was generated, to be compared with relatives of the missing persons. All missing persons but one had close living relatives, who were all found to be unrelated to the remains. The last person, who was born in Poland and had an Ashkenazi Jewish ancestry, had no close living relatives who survived the holocaust. His age when becoming missing was around 30 years, matching the age of death estimated for the skeletal remains.

Given the entire body of forensic evidence, the skeletal remains were determined to potentially belong to that person. However, the case was not closed, and the femur was left in storage for future investigation. A few years ago, it was discovered that the missing person has a distant (5^th^-degree) living relative, whose DNA was sampled as part of the investigation. The goal of this study was to generate genome-wide data for the unidentified femur. The data would be used to infer the ancestry of the genome. If confirmed as Ashkenazi Jewish, the genome would be compared to the known living relative of the missing person.

## Methods

### Approval

The study was approved by the relevant Israeli forensic authorities. The remains will be returned to the cave where it was found upon completion of the study. The single known distant relative of the missing person has been informed regarding the investigation and the manuscript.

### DNA extraction and sequencing

The femur was sent to Astrea Forensics (Santa Cruz, CA, USA) in July 2023 for DNA extraction and library preparation. DNA was extracted using the method of Rohland et al^28^. A second round of extraction was performed from the bone pellet of the original extraction. One library was prepared from each extraction using the Single Reaction Single-stranded LibrarY (SRSLY) method^29^. No UDG treatment was performed for correcting DNA damage.

Sequencing was performed on an Illumina MiSeq instrument, generating 76bp long paired-end reads. Sequencing of each of the two libraries generated about 1-2 million reads for the purpose of initial screening. The sequencing reads from the first two libraries were not used in downstream analyses.

For the purpose of the main analysis, a third library was generated using the same methods and sequenced to about 500 million reads.

### Genomic data processing

We trimmed and merged reads using AdapterRemoval v2^30^. We then mapped both merged and unmerged reads to the human reference genome (hs37d5) using bwa mem in single-end mode with default parameters^31^. We evaluated DNA damage using mapDamage^32^ and contamination using contamMix^33^. In the main analysis, we soft-clipped the reads to minimize residual damage biases using ATLAS^34^.

Given the extremely low coverage, and in order not to introduce any biases due to the use of specific reference panels, we did not attempt to impute missing variants^35^.

### Sex determination and uniparental markers

The sex of the individual was estimated by inspecting the number of reads mapping to the autosomes and each of the sex chromosomes (Figure 2).

The mitochondrial DNA haplogroup was inferred using Haplogrep 3^36^. The Y chromosome branch was inferred by YLeaf^37^, accepting single-read calls only when at least 90% of the bases agreed. We manually inspected variants defining branches upstream and downstream to the one returned by YLeaf to refine the branch assignment. The geographic distribution of the mitochondrial and Y haplogroups was assessed using FamilyTreeDNA Haplotree (familytreedna.com/public/y-dna-haplotree and familytreedna.com/public/mt-dna-haplotree) and Yfull (yfull.com/tree and yfull.com/mtree).

### Ancestry inference

We used principal components analysis (PCA) to infer the ancestry of the target genome. As reference genomes, we used 446 genomes of West Eurasian and North African populations from the Allen Ancient DNA Resource (AADR)^38^. We converted the target genome into “pseudo-haploid” using pileupCaller^39^ by sampling one read at random for each of the 597,573 SNPs in the AADR genomes. SNPs not covered by any read were left missing. We then used SmartPCA^40^ with the target genome projected on the axes formed by the AADR genomes. When using SmartPCA, least-squares projection was enabled and no outlier filtering iterations were applied.

We also inferred the ancestry of the target genome using ADMIXTURE^41^. We used the same set of reference genomes and pseudo-haploid target as in the PCA. We ran ADMIXTURE with default parameters and three values for the number of ancestral populations (K=6,7,8). To evaluate the robustness of the results, we ran ADMIXTURE a few times for each value of K, each time with a different seed.

## Results

### DNA extraction and sequencing

After DNA extraction and single-stranded library preparation, two libraries were sequenced for screening purposes with about 1-2 million reads each. The results showed poor DNA quality and widespread fragmentation, with only 4.5% and 3.4% of the reads mapping to the human genome, and mean fragment length (after merging paired reads) of 148 and 147 base pairs, in the two libraries, respectively.

Despite the low human DNA content, we decided to fully sequence the sample, resulting in 498.7 million reads. After removing duplicates, only 2.50 million (0.5%) mapped to the human genome, with a mean read length of 85bp. The mean human genome sequencing depth was 0.07x, and 5.8% of the genome was covered by any read. The mitochondrial DNA sequencing depth was 33x.

There was no evidence of contamination (using contamMix), with a maximum posteriori (MAP) contamination rate estimate of 0.00048 (2.5%–97.5% quantiles: 0.000077 to 0.0112). DNA damage was very minor, as apparent in a small increase in the frequency of C-to-T substitutions in read ends (Figure S1), as expected from a single-stranded library^42^.

### Sex and uniparental haplogroups

We inferred the sex of the individual based on the depth of sequencing across chromosomes (Figure 1). The results demonstrate nearly-uniform sequencing depth across the autosomes and between the X and Y chromosomes, suggesting that the genome belongs to a male individual, as inferred from the forensic investigation.

**Figure 1.**
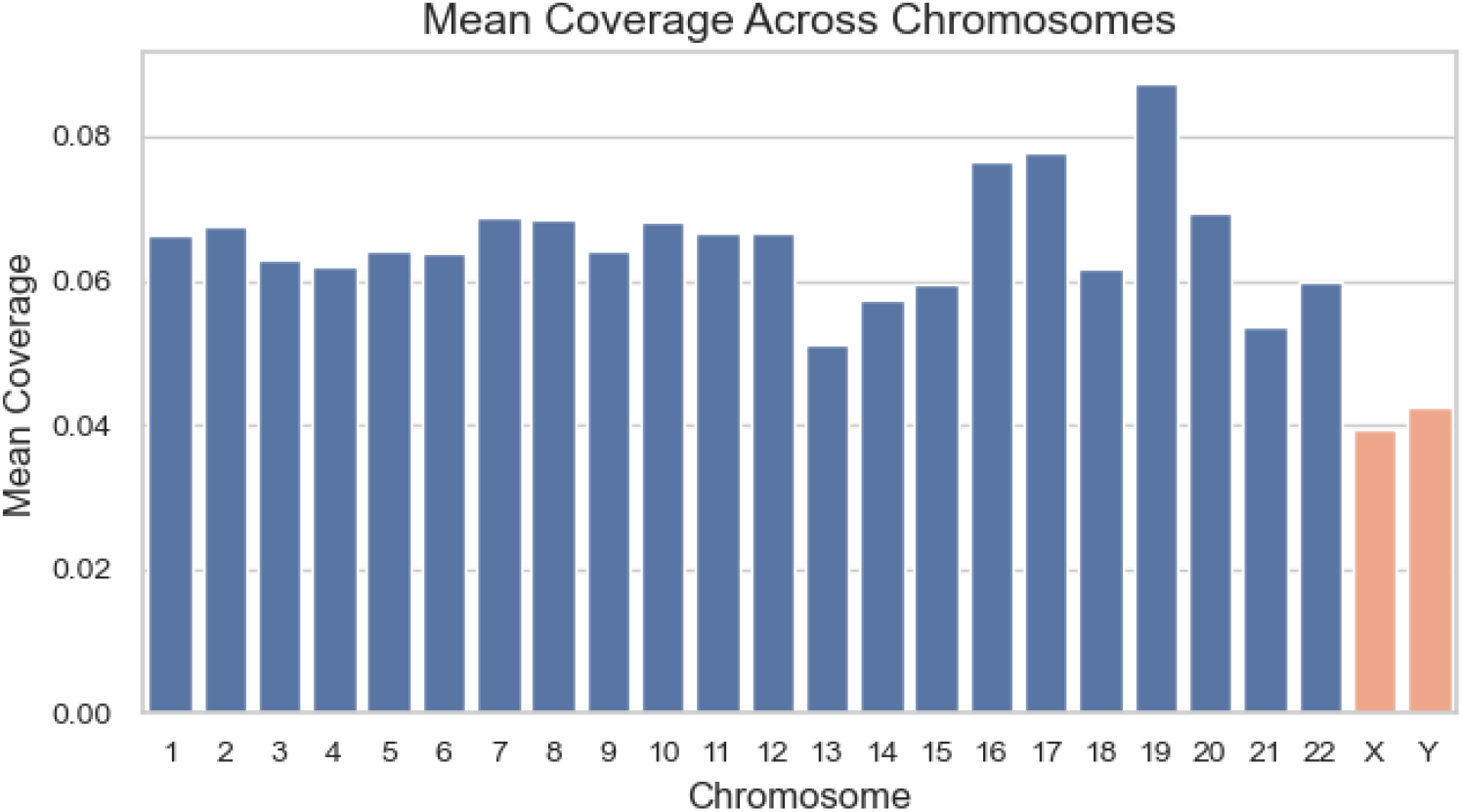
The mean sequencing depth across (nuclear) chromosomes.

The mitochondrial DNA haplogroup was T1a2, which is present across all of West Eurasia. The Y chromosome branch was G-FT276712 in YFull nomenclature (G-FT18229 in FTDNA nomenclature); the high missingness rates did not permit further refinement. G-FT276712 is present in current databases mostly in the Arabian Peninsula and the Levant.

### Genome-wide ancestry inference

We projected the target genome into PCA axes inferred from 446 reference genomes from West Eurasia and North Africa (Figure 2). The target genome clustered with Levant and North African populations: Egyptians, Bedouin, and Libyans. The number of SNPs used in the analysis was 36,098, which was shown to provide robust placement of genomes in PC space in the context of the Ashkenazi Jewish and Levant populations^43,44^. Thus, the genetic ancestry of the target genome is Arab, firmly excluding the hypothesis of an Ashkenazi Jewish origin.

**Figure 2.**
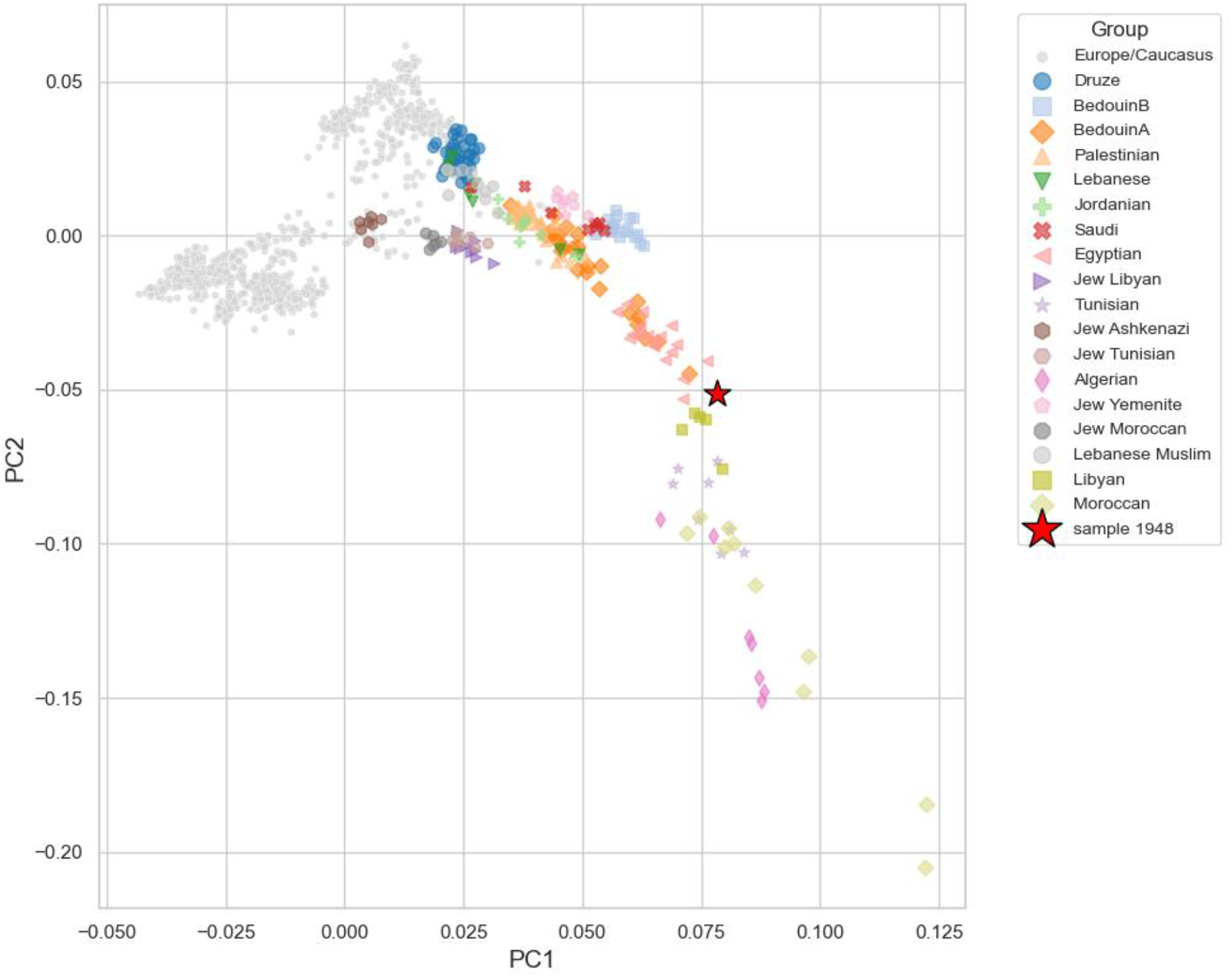
Projection of our sample on a PCA of modern reference populations from West Eurasia and North Africa. European and Caucasian individuals are plotted as small gray dots, while Middle-Eastern and North African individuals are plotted as colored circles. Our target sample is plotted as a large red star, clustering with Southern Levantine and North African populations.

To validate the ancestry assignment, we performed ancestry inference with ADMIXTURE. Focusing on K=6 ancestral components, we found that our target genome is inferred to have either North African or Middle Eastern ancestry, with the results varying across runs (Figure 3). We obtained similar results with K=7 and K=8 ancestral populations (Figure S2). Thus, we confirm that our target genome has no Ashkenazi Jewish genetic ancestry.

**Figure 3.**
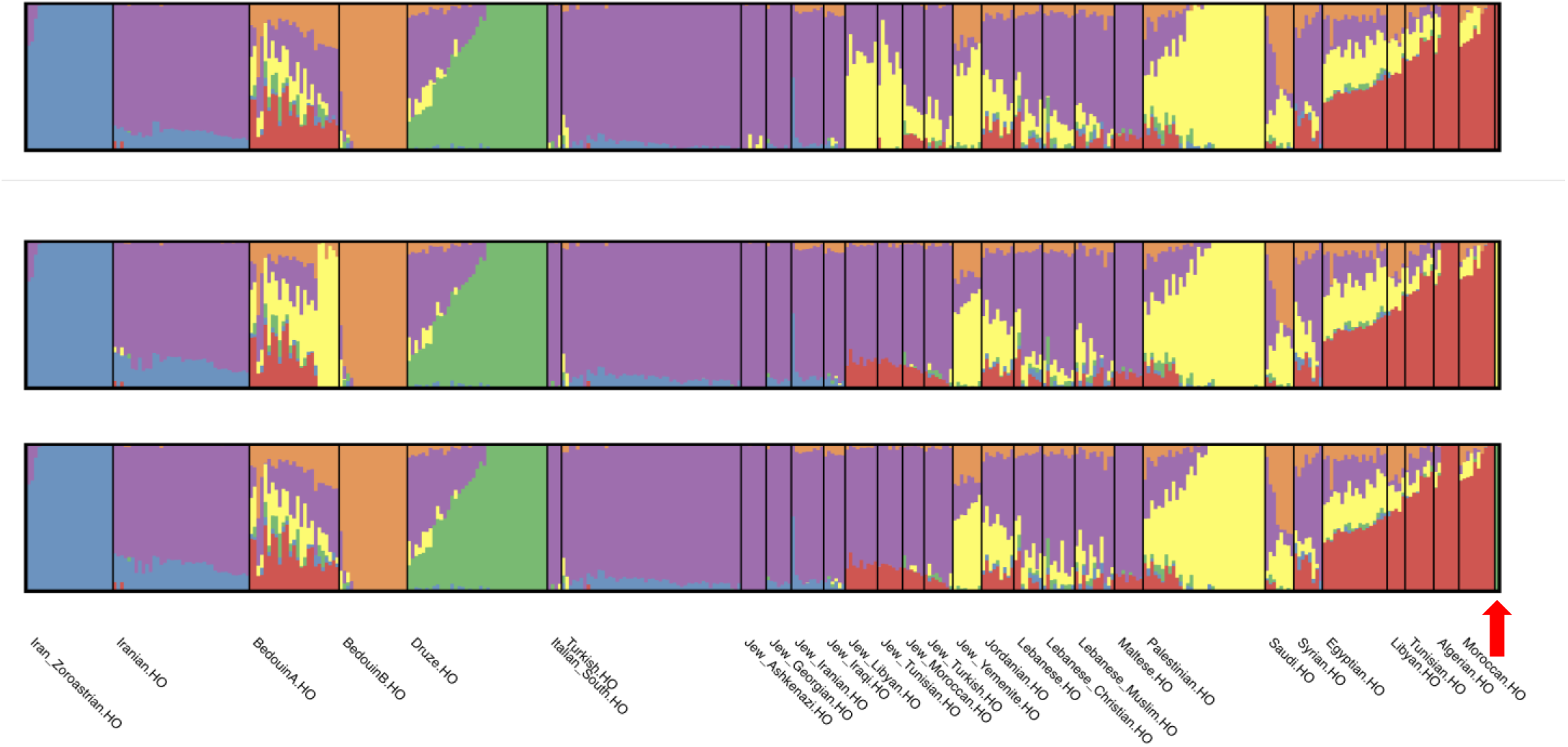
Ancestry proportions inferred by ADMIXTURE with K=6 ancestral population. We ran ADMIXTURE with the same set of West Eurasian and North African populations as used for the PCA. The figure only shows the results for the Middle Eastern and North African populations (HO: Human Origins dataset from AADR). Our target genome is highlighted with a red arrow. Each vertical bar corresponds to a single individual and is partitioned into K=6 colored segments representing the inferred ancestral populations. We show three panels corresponding to three different runs of ADMIXTURE, each time with a different seed for the random number generator. The ancestry of our target genome varies between being most similar to North African, Palestinian, and Druze.

## Discussion

To the best of our knowledge, ours is the first study to perform whole-genome sequencing on a middle-20^th^ century forensic sample from the Middle East. The results showed very poor DNA preservation. This is perhaps surprising given the success of previous ancient DNA studies from the region, including from prehistoric periods^44–51^, but may be due to the exposed state of the skeleton when detected.

The genetic ancestry of the bone was determined to be Arab, with results consistent between the autosomes and the Y chromosome and across methods. The highly missing data, as well as the underrepresentation of the Levant populations in reference datasets, did not permit higher resolution ancestry inference. For example, the ADMIXTURE analysis showed similar likelihood of North African and Palestinian ancestry. Nevertheless, the ancestry analysis convincingly excluded the possibility of an Ashkenazi Jewish ancestry. Therefore, the initial identification of the remains as belonging to a person of Ashkenazi origin has been refuted.

In conclusion, our results highlight the difficulty of extracting and sequencing DNA from middle-20^th^-century skeletal remains from the Levant. Nevertheless, the application of advanced experimental and computational methods from ancient DNA research enabled whole-genome sequencing (albeit sparse) of the sample, as well as ancestry assignment, providing critical investigative leads.

## Acknowledgements

We thank the DNA Doe Project and Kevin Lord for initial case management and bioinformatic analysis.

## Conflicts of interest

EV and SW are current or former employees and stock owners at MyHeritage. SC is a paid consultant and stock owner at MyHeritage. MyHeritage was not involved in the study.

## Funding

The study was supported by the forensic knowledge center grant (no. 1001584586) from the Israel Ministry of Innovation, Science & Technology to SC.

## Author contributions

Conceptualization: SC, JO; Formal Analysis: EV, SW, AL, ESB; Funding Acquisition: SC; Investigation: SC, EV, SW, AL, ESB, NA, BD, SA, YN, EM, NB, OK, DR, JO, RYL; Methodology: SC, RYL; Project Administration: SC, JO; Resources: JO; Software: EV, SW; Supervision: SC; Visualization: EV, SW, ESB, EM, SA; Writing – Original Draft Preparation: SC; Writing – Review & Editing: SC, EV, SW, RYL.

## Supplementary Figures

**Figure S1.**
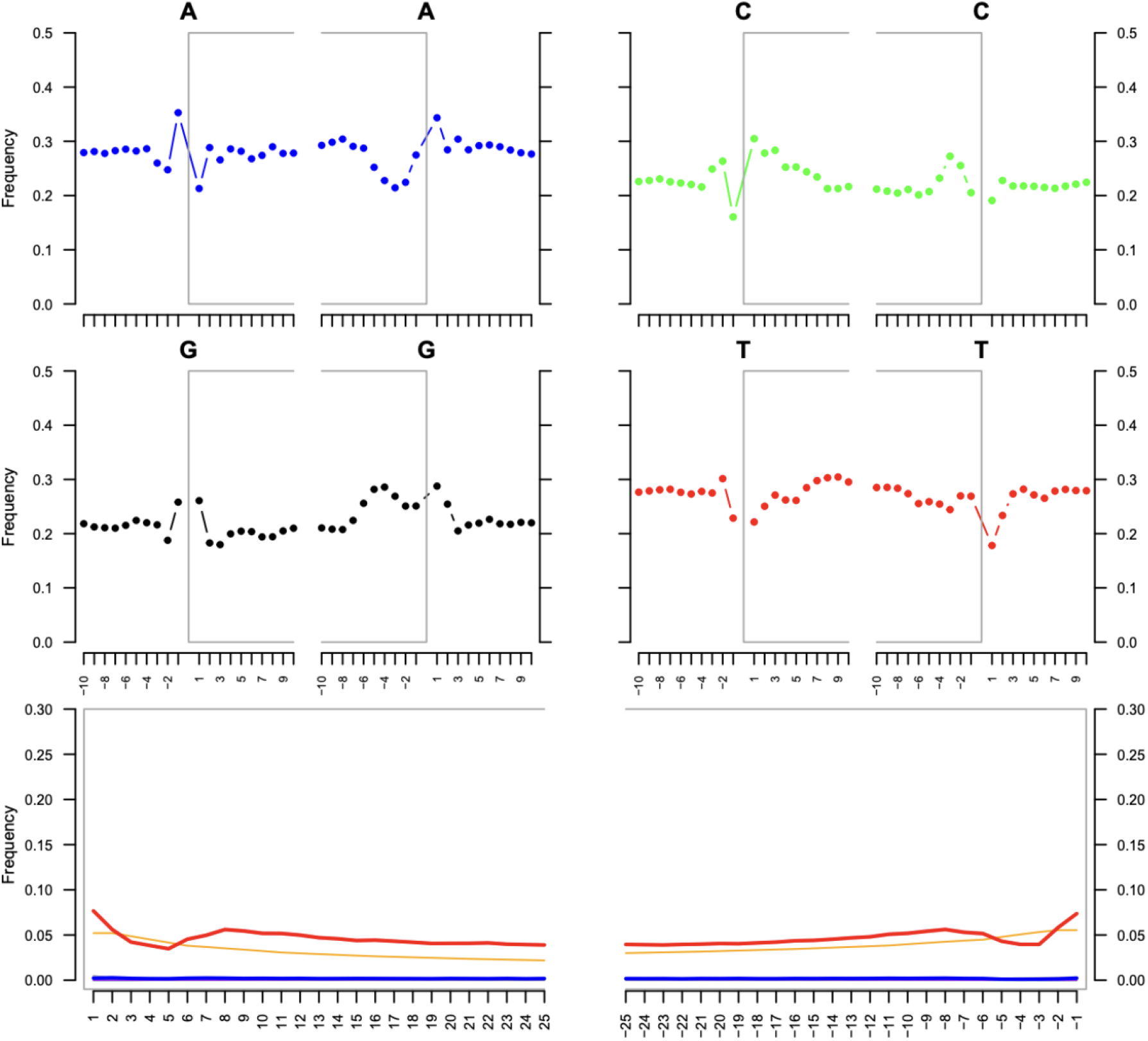
Nucleotide misincorporation patterns and fragment-end frequencies. The plots were generated using the mapDamage package, illustrating DNA damage patterns in ancient DNA samples. Top panels: base frequency outside and inside the read (the open grey box corresponds to the read). Bottom panel: position-specific substitutions from the 5’ (left) and the 3’ end (right). The following color codes were used: red: C → T; blue: G → A; orange: soft-clipped bases; purple: insertions relative to the reference.

**Figure S2.**
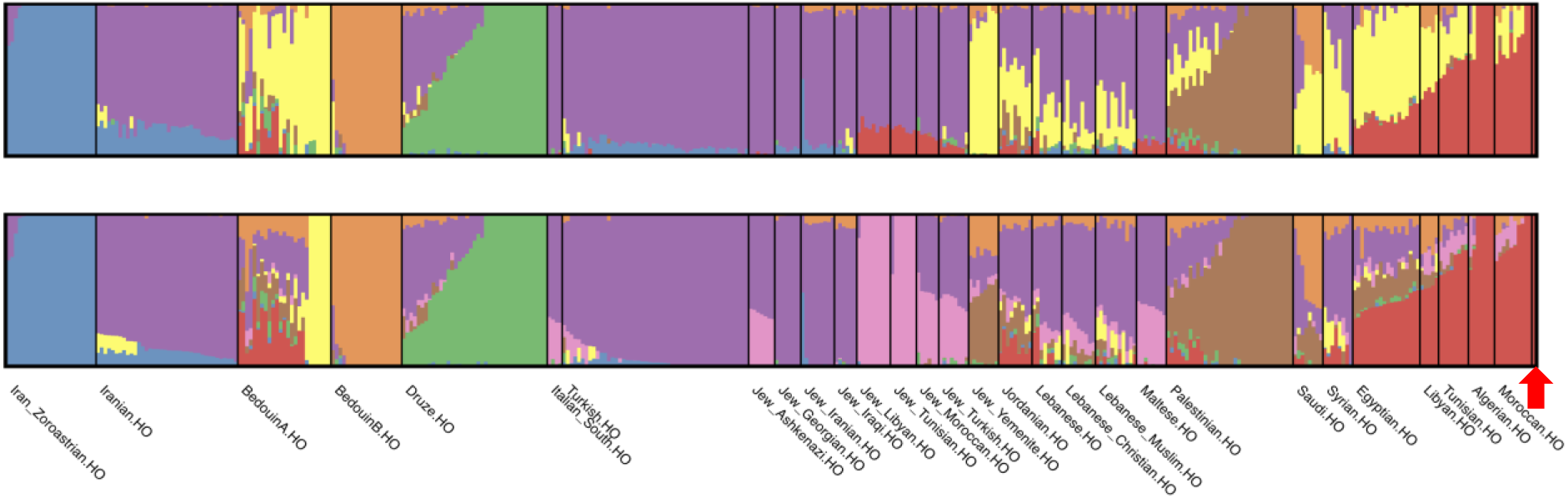
Replicate ADMIXTURE profiles for K=7 and K=8 ancestral populations. The figure was generated as in Figure 3, except that we used K=7,8 ancestral components. The target genome (red arrow) shows North African ancestry.

